# SlicerMorph Photogrammetry: An Open-Source Photogrammetry Workflow for Reconstructing 3D Models

**DOI:** 10.1101/2025.03.07.641939

**Authors:** Oshane O. Thomas, Chi Zhang, A. Murat Maga

## Abstract

**Context:** Accurate three-dimensional (3D) models of skeletal and other biological specimens are crucial for ecological and evolutionary research and teaching. Here, we present a streamlined, open-source workflow for 3D photogrammetry that allows researchers to construct 3D models using the 3D Slicer and the SlicerMorph ecosystem.

**Objectives and Methods:** We present an updated photogrammetry pipeline within the 3D Slicer ecosystem that combines advanced image masking via Segment Anything Model (SAM) with the open-source ODM reconstruction engine. Our approach systematically reduces artifacts in delicate cranial regions by automating background removal. We validate this pipeline against micro-CT references using multiple rodent skull specimens, quantifying accuracy through mean distance, root mean square error (RMSE), Hausdorff distance, and Chamfer distance.

**Results:** Our method consistently reduces average surface error by 10–15% compared to a previous open-source photogrammetry workflow. This gain is particularly pronounced around thin or fenestrated anatomical structures such as zygomatic arches, where the previous workflow created the most noise. Although occasional increases in Hausdorff distance reflect localized photography gaps, improved mean distance and RMSE underscore improved overall fidelity.

**Main Conclusions:** The 3D Slicer Photogrammetry extension provides a robust, fully open-source solution for generating high-quality 3D reconstructions from photographs. The streamlined pipeline lowers technical barriers to collecting reliable 3D data and fosters reproducible research across diverse biological collections.

## 1. Introduction

3D digitization has become a cornerstone in modern morphological research. It enables scientists to capture high-resolution data of skeletal remains, whole organisms, and other biological structures without physically disturbing the original specimens (Abhinav et al., 2021; Peng, 2008; Santana et al., 2019; Silva et al., 2023). By generating digital replicas, researchers can dissect complex forms for comparative analyses in functional morphology, taxonomic classification, biomechanics, and evolutionary biology (Irschick et al., 2022; Silva et al., 2023).

In particular, photogrammetry, which constructs a 3D surface model from multiple overlapping photographs taken at different positions, has experienced a surge in adoption due to its relatively low cost, portability, and ability to image objects of varied sizes. The resulting textured models capture both geometric and color information, facilitating a wide array of morphological and ecological interpretations. Furthermore, the ability to digitally archive specimens in high-fidelity 3D form is crucial for globally distributed research collections, especially when dealing with fragile or rare organisms (Blagoderov et al., 2012; Vollmar et al., 2010).

Despite the growing popularity of photogrammetry, many existing workflows rely on proprietary software that can be prohibitively expensive or impose licensing restrictions. In parallel, the open-source community has generated numerous structure-from-motion (SfM) tools, but these tools often require advanced scripting or specialized computing environments (Biegel, 2024; Patel et al., 2024; Vacca, 2019; Zhang & Maga, 2023).

A major bottleneck in many photogrammetry workflows is the need for background removal, which can be labor-intensive. Traditional approaches often require manual tracing or semi-automatic thresholding, introducing potential operator bias and risking the loss of fine structural details. Skull specimens with thin or fenestrated bony regions, such as the zygomatic arches of the skull, are especially susceptible to over-erosion during masking, resulting in incomplete or “broken” geometries (Zhang & Maga, 2023).

In addition, artifact removal (e.g., bridging across suture lines or “clogs” in narrow passages) can take a significant amount of time, limiting throughput for large sample sets. This is particularly problematic for studies that rely on subtle morphological signals to distinguish closely related groups.

Dashti et al., 2022 suggest that discrepancies as small as 0.1–0.2 mm may obscure functional traits or early signals of morphological divergence in small mammals . These small-scale differences can reflect adaptive changes in dentition, cranial morphology, or muscle attachment sites.

The study by Zhang & Maga (2023) showed the possibility of consolidating various open-source tools into a pipeline for generating 3D textured models from photographs using a low-cost, portable setup. This initial attempt was not quite streamlined and required manual background removal and semi-automatic masking that was only partially effective, which resulted in reduced fidelity, artifacts, and noise in geometrically complex regions such as the pterygoid areas and zygomatic processes of the cranium. Nonetheless, the study demonstrated the feasibility of using low-cost commodity hardware and open-source to acquire a large number of photographs and build 3D textured models that are sufficiently accurate for most morphometric studies or teaching purposes.

Here, we improve on this study by integrating the Segment Anything Model (SAM) (Kirillov et al., 2023) with the structure-from-motion reconstruction (NodeODM, part of OpenDroneMap project (Patel et al., 2024)) and combining these tools within the 3D Slicer (Fedorov et al., 2012) environment to unify data acquisition, segmentation, reconstruction, and final model generation. The updated pipeline aims to:

1. Streamline background masking while improving the quality of the obtained mask,
2. Explore new parameters introduced to ODM since the publication of the previous study to improve 3D model generation,
3. Overall, it simplifies user interaction by minimizing manual operator input.

The SlicerMorph extension, developed within the 3D Slicer ecosystem, allows users to seamlessly continue their workflow, whether for visualization, landmarking or other sorts of morphometric analysis (Rolfe et al., 2021).

In the next section, we quantitatively assess the improvements to the geometric accuracy of the models reconstructed from the Photogrammetry extension by comparing the models generated from both workflows to their ground-truth model that was acquired by high-resolution micro-CT. We provide quantitative (mean distance, RMSE, Hausdorff, Chamfer) and qualitative (visual) evidence of improved capture of intricate skeletal elements.

## 2. Materials and Methods

### 2.1. Overview of the Extension and Workflow

The workflow begins with importing photographs (e.g., skulls or other biological specimens) into the Photogrammetry module, which leverages SAM to isolate specimens from image backgrounds rapidly. Masking can be performed batch-wise or refined on an individual-image basis, significantly reducing the manual effort typically required. Following image preparation, the extension directly integrates with NodeODM to reconstruct high-resolution 3D models, which otherwise may require commercial software products. The extension extends the existing open-source SlicerMorph ecosystem in 3D Slicer for 3D morphology and morphometrics analysis. It facilitates publication-quality 3D model reconstructions, enabling ecologists and evolutionary biologists to quantify morphological variation readily, archive museum collections, or share models openly.

### 2.2. Software Architecture and Dependencies

Photogrammetry was developed for 3D Slicer (version 5.8; (Fedorov et al., 2012)) and is currently available through the Slicer Extension Manager for the Linux operating system. It depends on two primary Slicer extensions—PyTorch and SlicerMorph—as well as several Python libraries:

○ **segment-anything** for deep-learning-based image masking (Kirillov et al., 2023).
○ **pyodm** for interacting with OpenDroneMap’s NodeODM (Patel et al., 2024).
○ **OpenCV** and **Pillow** for image processing and I/O.

NodeODM reconstruction requires Docker and, optionally, the NVIDIA Container Toolkit for GPU acceleration for NVIDIA GPUs, for which specific guidelines for how to set these environments can be found on the web as they can differ between various Linux operating systems. For users who do not want to deal with the technical aspects of setting up containers or users on Windows and Mac operating systems, we recommend using the free MorphoCloud On-Demand platform, in which all necessary libraries are pre-configured, to avoid extensive local setup. MorphoCloud On-Demand instances provide a GPU-equipped, turnkey, powerful cloud environment with dozens of cores and over 100GB of RAM. This enables users to access the entire pipeline that scales up to thousands of photographs without hitting hardware restrictions.

### 2.3. Installation and Setup

We describe two primary installation options for the Photogrammetry Extension for 3D Slicer:

#### 2.3.1. MorphoCloud On Demand (Recommended)

○ Users access the MorphoCloud platform (https://github.com/MorphoCloud/MorphoCloudInstances), a GPU-enabled environment preconfigured with all required software (Docker, NodeODM, NVIDIA Container Toolkit, PyTorch).

○ Launch the Photogrammetry module directly from the SlicerMorph submenu of the 3D Slicer’s Module Selector to begin processing immediately.

#### 2.3.2. Local Installation On Linux

○ Install Docker, NVIDIA GPU drivers, and the NVIDIA Container Toolkit if GPU acceleration is desired.
○ Confirm Docker functionality (e.g., **docker run --rm --gpus all nvidia/cuda:11.8.0-base**).
○ Install 3D Slicer (version 5.8) and use the Extension Manager (Linux) to install “SlicerMorph Photogrammetry. “ PyTorch and SlicerMorph will be automatically installed as dependencies.
○ On Windows and macOS, users must manually clone and load the repository as a scripted module (more detailed instructions are provided online).

### 2.4. Image Acquisition and Masking

While robust to various image acquisition methods, we recommend consistent illumination and minimal background clutter for optimal results, following the recommendations in Zhang and Maga (2023). Images can be captured using a basic DSLR camera on a programmable turntable within a diffused lighting setup. For effective reconstruction, photographs should have approximately 70–80% overlap.

Once imported into the extension, the images are masked using SAM. Depending on hardware capability, users select the masking resolution (full, half, or quarter). Low-resolution masking is computationally less taxing and faster at the expense of accuracy. Two masking methods are available:

○ <u>Batch Masking:</u> In one representative image, a bounding box is placed around the specimen, and it is propagated automatically to all images.
○ <u>Single Image Masking:</u> Individual adjustments are made to refine challenging images, including placing inclusion/exclusion markers for precision.

Masked images and binary masks, preserving essential camera metadata, are automatically stored for subsequent 3D reconstruction steps.

### 2.5. Photogrammetry Reconstruction via NodeODM

After image masking, users reconstruct 3D meshes through NodeODM, accessible directly from the Photogrammetry extension:

#### 2.5.1. Optional Scaling (via ArUco Markers)

Users may include ArUco markers in images to provide accurate physical scaling of the specimen. The extension automates the creation of Ground Control Point (GCP) files (“Find-GCP”), allowing NodeODM to incorporate scale and position accuracy into the final model. For instructions on how to create a local coordinate system using Aruco markers, see Section 2.3 of the Supplemental Online Material from Zhang et al., 2023.

#### 2.5.2. Launching NodeODM

The module provides a simple interface for launching a local NodeODM container via Docker or connecting to an existing remote server. The user should provide the IP address and the port number NodeODM is running for the latter.

#### 2.5.3. Reconstruction Parameters

The module offers recommended settings, balancing detail with computational efficiency for reconstructing on Morphocloud On-Demand instances for most morphological applications:

○ mesh-octree-depth = 12
○ mesh-size = 300,000–500,000
○ feature-quality = ultra
○ pc-quality = high
○ no-gpu = false (recommended for stable texturing)

Users initiate reconstruction via the “Run NodeODM Task” button, and the module provides real-time updates.

#### 2.5.4. Model Retrieval

Completed 3D models (OBJ and associated texture images) are automatically downloaded, organized, and available for direct import into 3D Slicer for immediate visualization and measurement. The reconstruction parameters and configuration details are saved as JSON files to facilitate reproducibility.

### 2.6. Worked Example: Mountain Beaver Skull (UWBM 82409)

To illustrate the workflow, we reconstructed a mountain beaver (*Aplodontia rufa*) skull (specimen UWBM 82409) from 320 images. Following Zhang & Maga (2023), images were taken to ensure high overlap (∼70–80%) and consistent illumination. After batch-masking the images in about 20 minutes, we refined select images manually, for which automatic removal of the mounting platform and clay had failed.

NodeODM is launched locally (GPU enabled) with the default parameters described above, and the user optionally provides a GCP list for correct physical scaling of the specimen. Reconstruction yielded a detailed, textured mesh that was automatically imported back into 3D Slicer for evaluation. The complete set of images, masked outputs, and final reconstructed models are available publicly at https://app.box.com/shared/static/z8pypqqmel8pv4mp5k01philfrqep8xm.zip.

A graphical overview of this example is presented in Figure 1, illustrating the masking and reconstruction pipeline.

**Figure 1.**
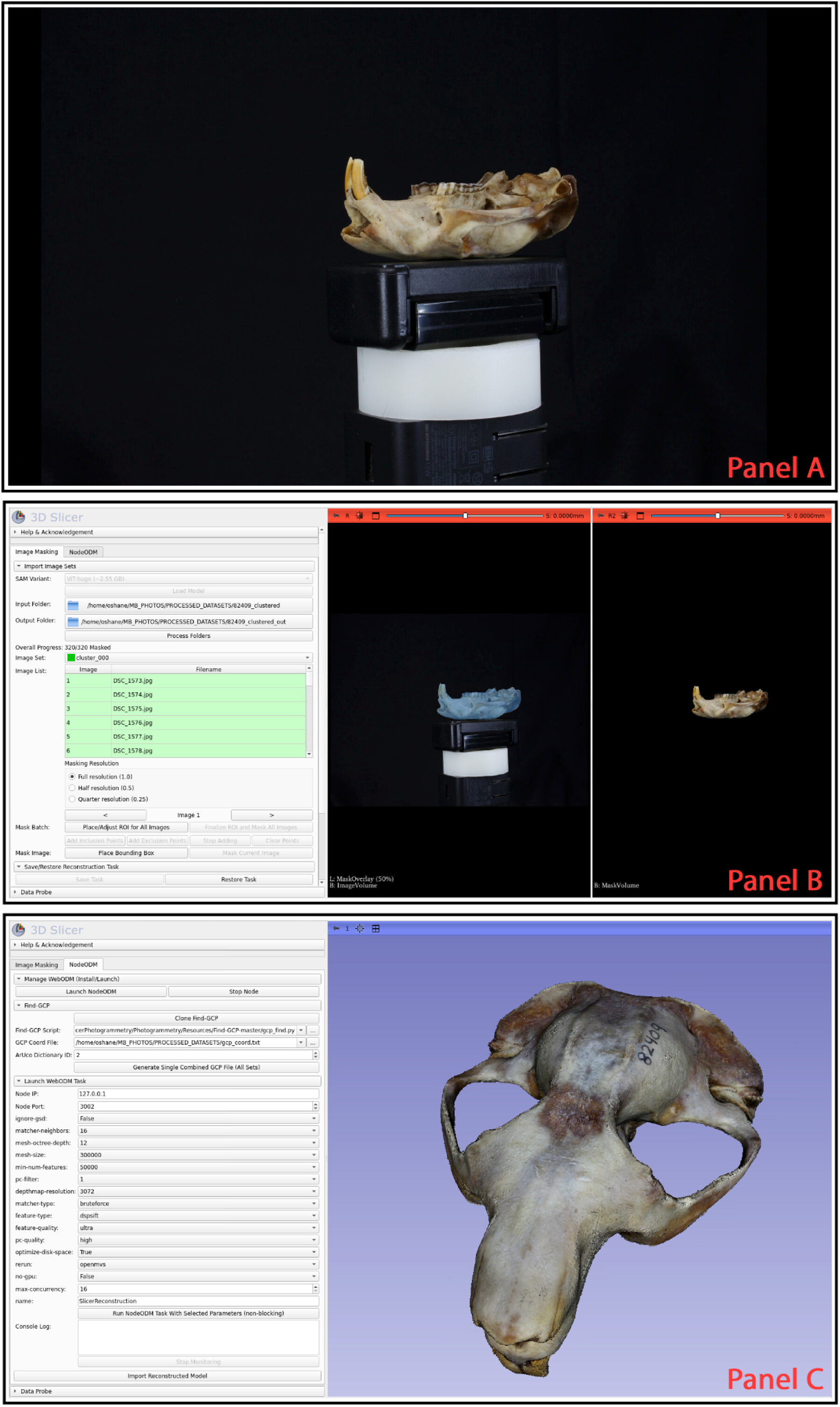
Overview of the Photogrammetry Extension workflow using a mountain beaver skull dataset. (1A) Example of one original photograph from the 320-image dataset, which was taken as the specimen mounted on a turntable. The Photogrammetry user interface consists of two module tabs (Image Masking and NodeODM) that focus on different aspects of the workflow. (1B) The Image Masking interface displaying a fully masked dataset. The left 2D view shows the original photograph overlaid with a blue mask generated using the Segment Anything Model, highlighting the segmented specimen. The right 2D view presents the same image post-segmentation, with the background removed, isolating the skull. (1C) ODM interface post-photogrammetry reconstruction with default parameters set in the Find GCP and Launch NodeODM Task subsections. The resulting reconstructed and textured 3D skull model is displayed imported directly into the 3D viewport via the ’Import Reconstructed Model’ functionality.

### 2.7. Optimizing Photogrammetry Parameters via Taguchi Design

We determined the optimal NodeODM reconstruction settings for the *Aplodontia rufa* skulls dataset through a Taguchi L16 design (Montgomery, 2017), varying seven major parameters such as mesh-octree-depth, mesh-size, and ignore-gsd while holding other baseline factors constant (e.g., orthophoto-resolution, texturing-single-material).

We programmatically submitted masked images (plus optional ground-control points) to a local NodeODM instance using the Python pyodm library to facilitate this process.

This approach systematically enumerated 16 unique parameter combinations (e.g., matcher-neighbors = 0 or 16, pc-filter = 1, 2, or 3) and recorded the resulting textured models.

Because slight misalignments can persist, we first applied a 3-point registration to each reconstructed mesh to approximate alignment with a micro-CT “gold standard” scan. We then refined each alignment via Iterative Closest Point (ICP) using Open3D (Zhou et al., 2018). It should be noted that these alignments are “rigid” in nature and have no effect on the geometry of the reconstructed models; they simply alter the position of the model in 3D space. Reconstructed and reference meshes were uniformly sampled to point clouds (260,000 points) for distance-based error calculations (mean, RMSE, Hausdorff, and standard deviation). Finally, we ranked parameter combinations by mean distance from reconstruction to micro-CT (lowest is best). This Taguchi-based methodology allowed us to evaluate a range of parameters efficiently while balancing runtime and resource considerations. The best configuration—listed in

Table 2—provided minimal geometric error across multiple metrics.

### 2.8. Code and Data Availability

The code, scripts, and user documentation for the Photogrammetry extension are hosted at https://github.com/SlicerMorph/SlicerPhotogrammetry and permanently archived via Zenodo with a DOI.

The example dataset (specimen UWBM 82409) is provided at https://app.box.com/shared/static/z8pypqqmel8pv4mp5k01philfrqep8xm.zip and is publicly available on OSF [or a similar repository] under a permanent DOI. The repository also includes complete pipeline configurations (JSON files), enabling users to replicate our procedures fully.

### 2.9. Use of Large Language Models or AI

During manuscript and software development, the authors used ChatGPT (version 4.0, “Pro” mode) as an assistant for Python scripting, debugging, manuscript proofreading, and improving clarity and readability. All AI-generated code was manually verified for correctness and license compliance, and all AI-assisted text was thoroughly revised by authors for scientific accuracy and journal compliance.

### 2.10. Summary of Reproducibility

The described photogrammetry pipeline integrates entirely open-source tools and standardized protocols, enabling robust reproducibility. Detailed instructions, automated masking, explicit software dependencies, and openly archived code/data ensure that researchers can replicate our results across diverse morphological datasets with minimal setup and without proprietary software restrictions.

## 3. Results/Discussion

### 3.1. Overview of Model Quality and Accuracy

Across the *Aplodontia rufa* skull dataset, the updated Photogrammetry workflow systematically yielded more accurate 3D models than the previous pipeline (Zhang & Maga, 2023). Across 14 specimens, we observed a ∼10–15% decrease in mean surface error relative to the micro-CT reference (Table 1, Fig. 2). This improvement was particularly evident in delicate bony structures (e.g., zygomatic arches, orbital rims), where the older approach often generated small artifacts or over-smoothed regions. By contrast, the new method’s integration of the SAM and refined ODM parameters produced smoother surfaces with fewer extraneous polygons.

**Table 1.**
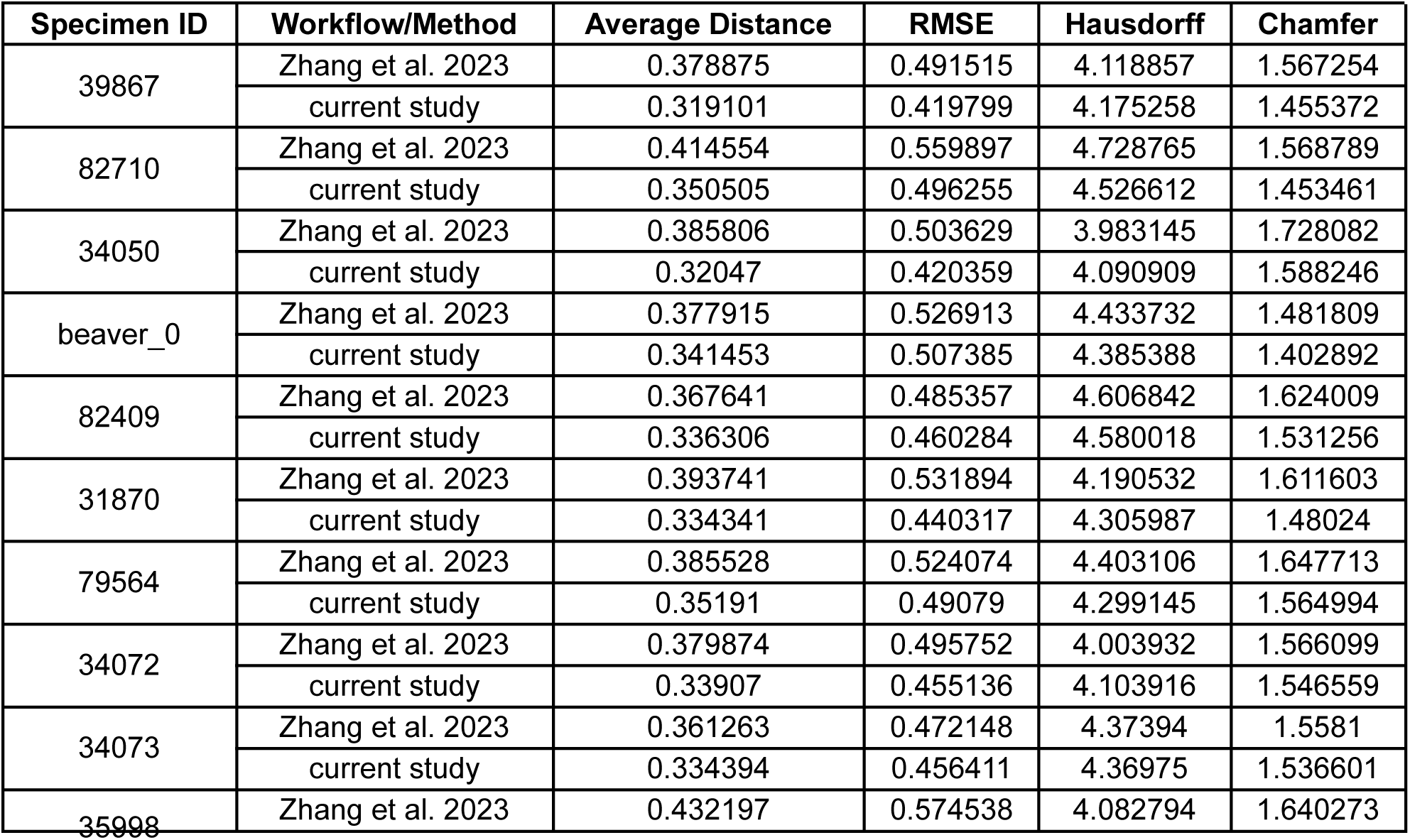
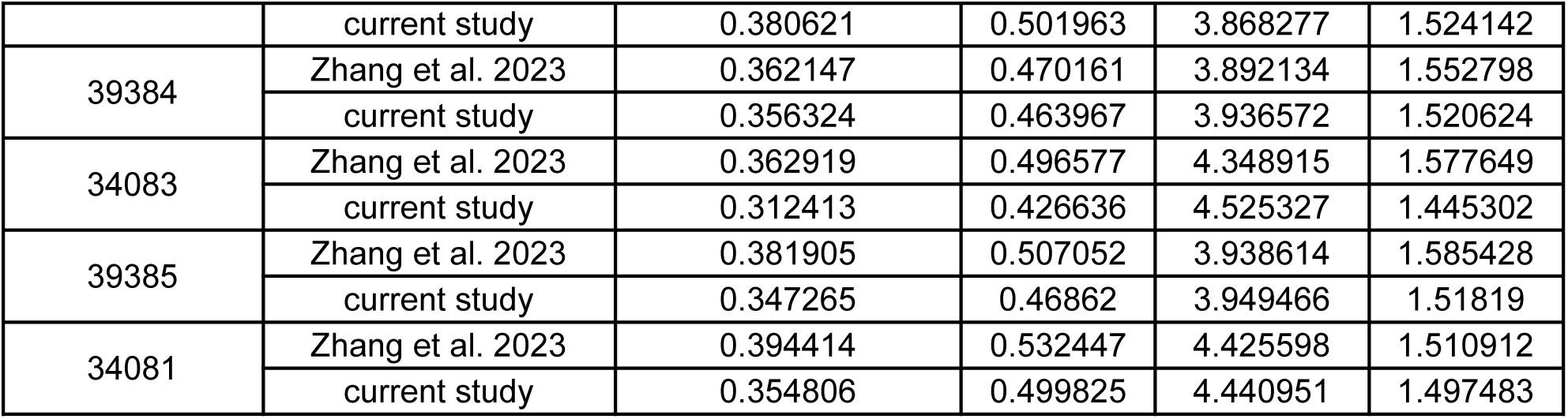
Reconstruction errors for two photogrammetry workflows (Zhang et al. 2023 vs. the updated workflow) compared to micro-CT references. Models were aligned to the CT via ICP, then uniformly sampled to measure Euclidean (L2) distances, RMSE (emphasizing outliers), Hausdorff distance (worst local discrepancy), and Chamfer distance (bidirectional mismatch). Across specimens, the new workflow generally yields lower average and RMSE values, with Hausdorff and Chamfer distances remaining comparable or improved. “UWBM” indicates specimens from the University of Washington Burke Museum; “beaver0” is from a private collection.

| Specimen ID | Workflow/Method | Average Distance | RMSE | Hausdorff | Chamfer |
| --- | --- | --- | --- | --- | --- |
| 39867 | Zhang et al. 2023 | 0.378875 | 0.491515 | 4.118857 | 1.567254 |
|  | current study | 0.319101 | 0.419799 | 4.175258 | 1.455372 |
| 82710 | Zhang et al. 2023 | 0.414554 | 0.559897 | 4.728765 | 1.568789 |
|  | current study | 0.350505 | 0.496255 | 4.526612 | 1.453461 |
| 34050 | Zhang et al. 2023 | 0.385806 | 0.503629 | 3.983145 | 1.728082 |
|  | current study | 0.32047 | 0.420359 | 4.090909 | 1.588246 |
| beaver_0 | Zhang et al. 2023 | 0.377915 | 0.526913 | 4.433732 | 1.481809 |
|  | current study | 0.341453 | 0.507385 | 4.385388 | 1.402892 |
| 82409 | Zhang et al. 2023 | 0.367641 | 0.485357 | 4.606842 | 1.624009 |
|  | current study | 0.336306 | 0.460284 | 4.580018 | 1.531256 |
| 31870 | Zhang et al. 2023 | 0.393741 | 0.531894 | 4.190532 | 1.611603 |
|  | current study | 0.334341 | 0.440317 | 4.305987 | 1.48024 |
| 79564 | Zhang et al. 2023 | 0.385528 | 0.524074 | 4.403106 | 1.647713 |
|  | current study | 0.35191 | 0.49079 | 4.299145 | 1.564994 |
| 34072 | Zhang et al. 2023 | 0.379874 | 0.495752 | 4.003932 | 1.566099 |
|  | current study | 0.33907 | 0.455136 | 4.103916 | 1.546559 |
| 34073 | Zhang et al. 2023 | 0.361263 | 0.472148 | 4.37394 | 1.5581 |
|  | current study | 0.334394 | 0.456411 | 4.36975 | 1.536601 |
| 35998 | Zhang et al. 2023 | 0.432197 | 0.574538 | 4.082794 | 1.640273 |
|  | current study | 0.380621 | 0.501963 | 3.868277 | 1.524142 |
| 39384 | Zhang et al. 2023 | 0.362147 | 0.470161 | 3.892134 | 1.552798 |
|  | current study | 0.356324 | 0.463967 | 3.936572 | 1.520624 |
| 34083 | Zhang et al. 2023 | 0.362919 | 0.496577 | 4.348915 | 1.577649 |
|  | current study | 0.312413 | 0.426636 | 4.525327 | 1.445302 |
| 39385 | Zhang et al. 2023 | 0.381905 | 0.507052 | 3.938614 | 1.585428 |
|  | current study | 0.347265 | 0.46862 | 3.949466 | 1.51819 |
| 34081 | Zhang et al. 2023 | 0.394414 | 0.532447 | 4.425598 | 1.510912 |
|  | current study | 0.354806 | 0.499825 | 4.440951 | 1.497483 |

**Table 2.** Top-performing Taguchi L16 configuration for NodeODM reconstruction, along with unidirectional error metrics relative to micro-CT scans. All distances are in millimeters. “E → G” indicates distances from Experimental reconstruction to Ground truth; “G → E” indicates the reverse. Differences in E → G and G → E arise when the micro-CT includes internal geometry not captured by surface-based photogrammetry, thereby inflating the G → E distances when the reference points lack corresponding structures in the reconstruction.

| Parameter/Metric | Value |
| --- | --- |
| ignore-gsd | FALSE |
| matcher-neighbors | 16 |
| mesh-octree-depth | 12 |
| mesh-size | 300 000 |
| min-num-features | 50 000 |
| pc-filter | 2 |
| feature-quality | ultra |
| pc-quality | ultra |
| mean_dist_E→G | 0.300535 |
| rmse_E→G | 0.402543 |
| hausdorff_E→G | 3.918398 |
| std_E→G | 0.267805 |
| mean_dist_G→E | 1.064301 |
| rmse_G→E | 1.550204 |
| hausdorff_G→E | 8.31234 |
| std_G→E | 1.127119 |
| chamfer | 1.364836 |

**Figure 2.**
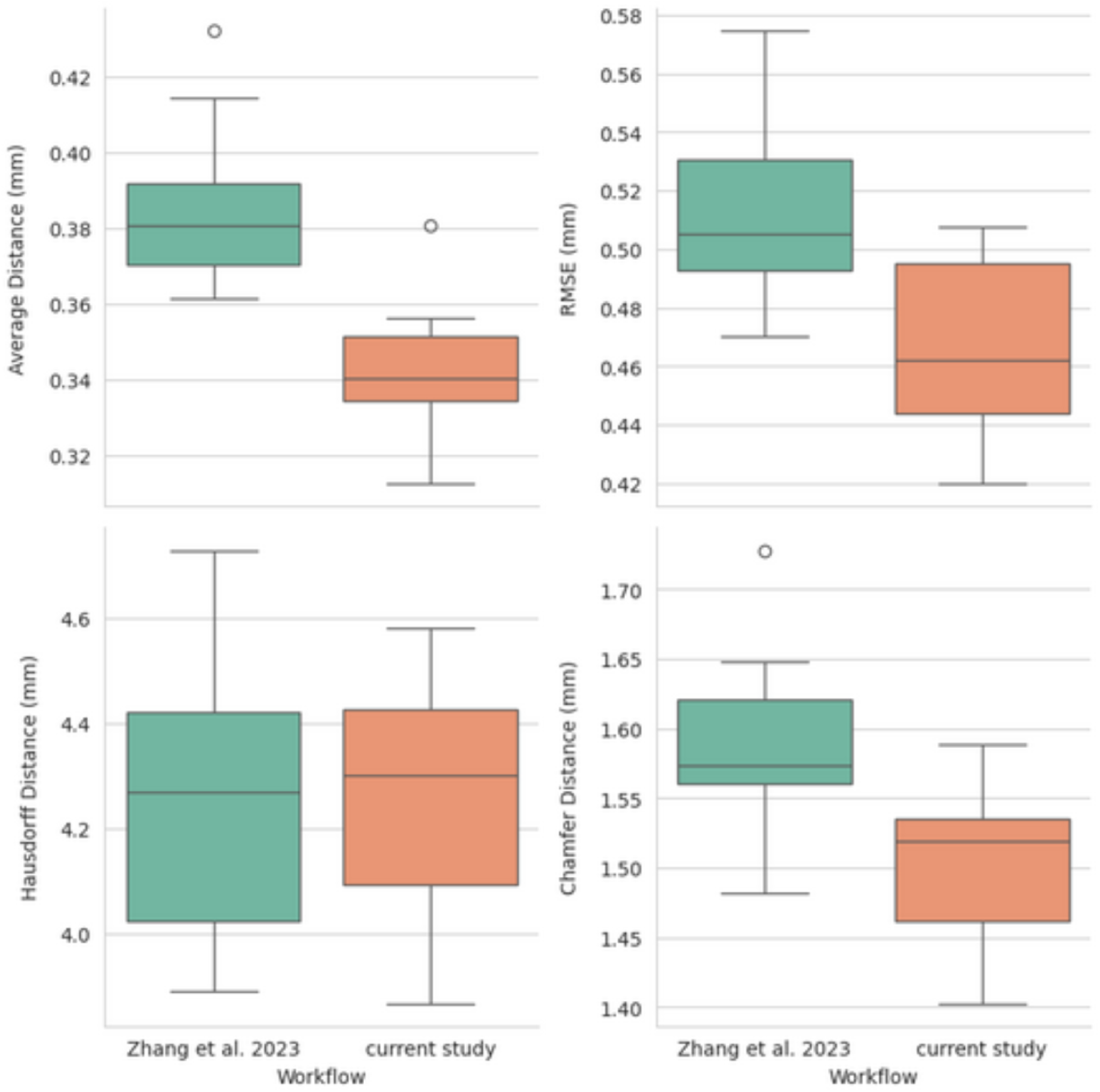
Box-plot comparison of reconstruction errors from Zhang et al. (2023) versus our updated Photogrammetry workflow, using micro-CT data as the ground-truth reference. Each panel shows one of four error metrics—Average Distance, RMSE, Hausdorff Distance, and Chamfer Distance—across multiple rodent skull specimens. The updated workflow yields consistently lower Average Distance and RMSE values (by about 10–15%) in most specimens, indicating improved overall fidelity in delicate cranial regions. Although Hausdorff and Chamfer distances vary by specimen—occasionally reflecting incomplete photo coverage—the overall trend suggests the new pipeline provides more accurate and reproducible 3D models for ecological and evolutionary research.

Alongside these quantitative gains in average distance and root mean square error (RMSE), we noted qualitatively higher fidelity (Fig. 3). Zygomatic arches and foramina, often prone to background noise or partial mis-segmentation in the previous study, were reconstructed with fewer breaks or bridging polygons. Although Hausdorff and Chamfer distance sometimes varied by specimen—chiefly when portions of the skull were incompletely photographed—global mean error and RMSE improvements indicate a more consistent capture of external morphology.

**Figure 3.**
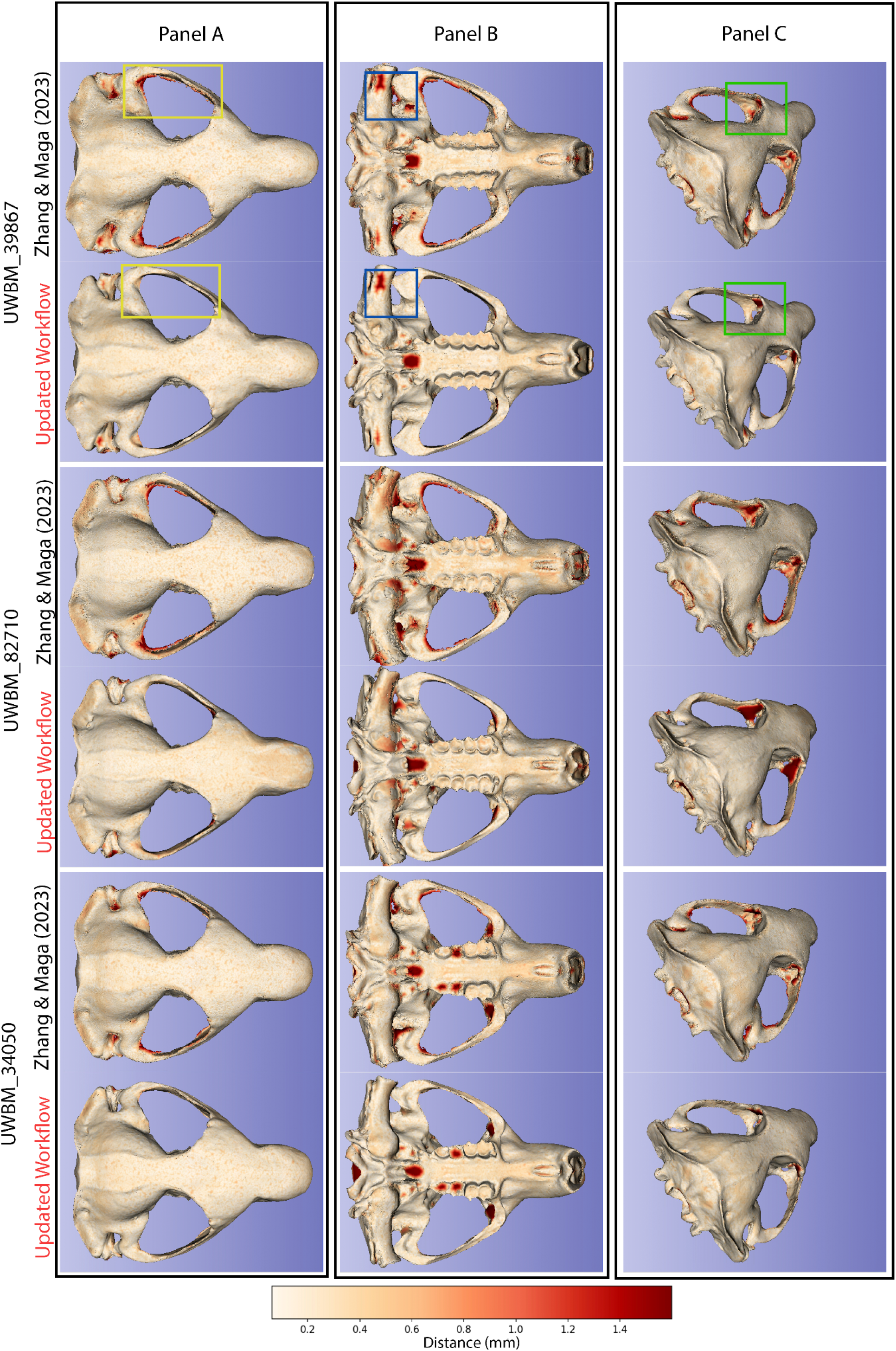
Side-by-side photogrammetric reconstructions of three *Aplodontia rufa* (mountain beaver) skull specimens—UWBM 39867 (top two rows), UWBM 82710 (middle two rows), and UWBM 34050 (bottom two rows)—shown in A (dorsal view), B (ventral view), and C (oblique posterior-lateral view). For each specimen and orientation, the upper image is derived from the Zhang & Maga (2023) workflow, and the lower image is reconstructed via the Photogrammetry extension being presented. Warmer colors in the distance maps (bottom scale in millimeters) indicate larger deviations, whereas cooler colors signify closer agreement. Across all three specimens, the new workflow yields fewer triangular artifacts around the orbital and zygomatic boundaries avoids clogging critical foramina, and reduces over-smoothed patches in under-photographed areas. Its enhanced Segment Anything Model–based masking more accurately separates the specimen from the background, resulting in consistently improved model fidelity and more anatomically faithful reconstructions.

### 3.2. Comparisons Between Old and New Workflows

Compared to the previous study, the updated pipeline consistently reduced unidirectional discrepancy from (E)xperimental reconstructed mesh to (G)round-truth mesh (E → G) by 0.04–0.07 mm, allowing finer capture of delicate bone edges. For example, in specimen UWBM 82710, the average error dropped from 0.4146 mm to 0.3505 mm, while RMSE fell from 0.5599 mm to 0.4963 mm. These gains are not merely technical but translate into fewer hours of manual editing since morphological landmarks now appear more sharply defined. Although Hausdorff’s distance occasionally increased slightly (e.g., from ∼4.37 mm to ∼4.53 mm) in some specimens, this typically reflected localized deficits, such as reflective areas or thin edges missed by the camera. Because ecological analyses often prioritize overall shape integrity over single outliers, the strong improvements in mean and RMSE are likely of greater practical importance.

Visual assessments reinforce these numeric trends. Due to suboptimal image masking, the prior pipeline occasionally “clogged” the pterygoid or palatal regions with extraneous mesh. In contrast, our approach better delineated these structures (Fig. 3). Warmer colors in deviation maps remain largely restricted to the periphery, indicating that interior cranial regions are aligned well with micro-CT references. The consistent preservation of fine anatomical details, especially around delicate areas such as the zygomatic arches and orbital margins, demonstrates the improved fidelity and reduced artifacts achieved by our updated photogrammetry workflow (Fig. 4).

**Figure 4.**
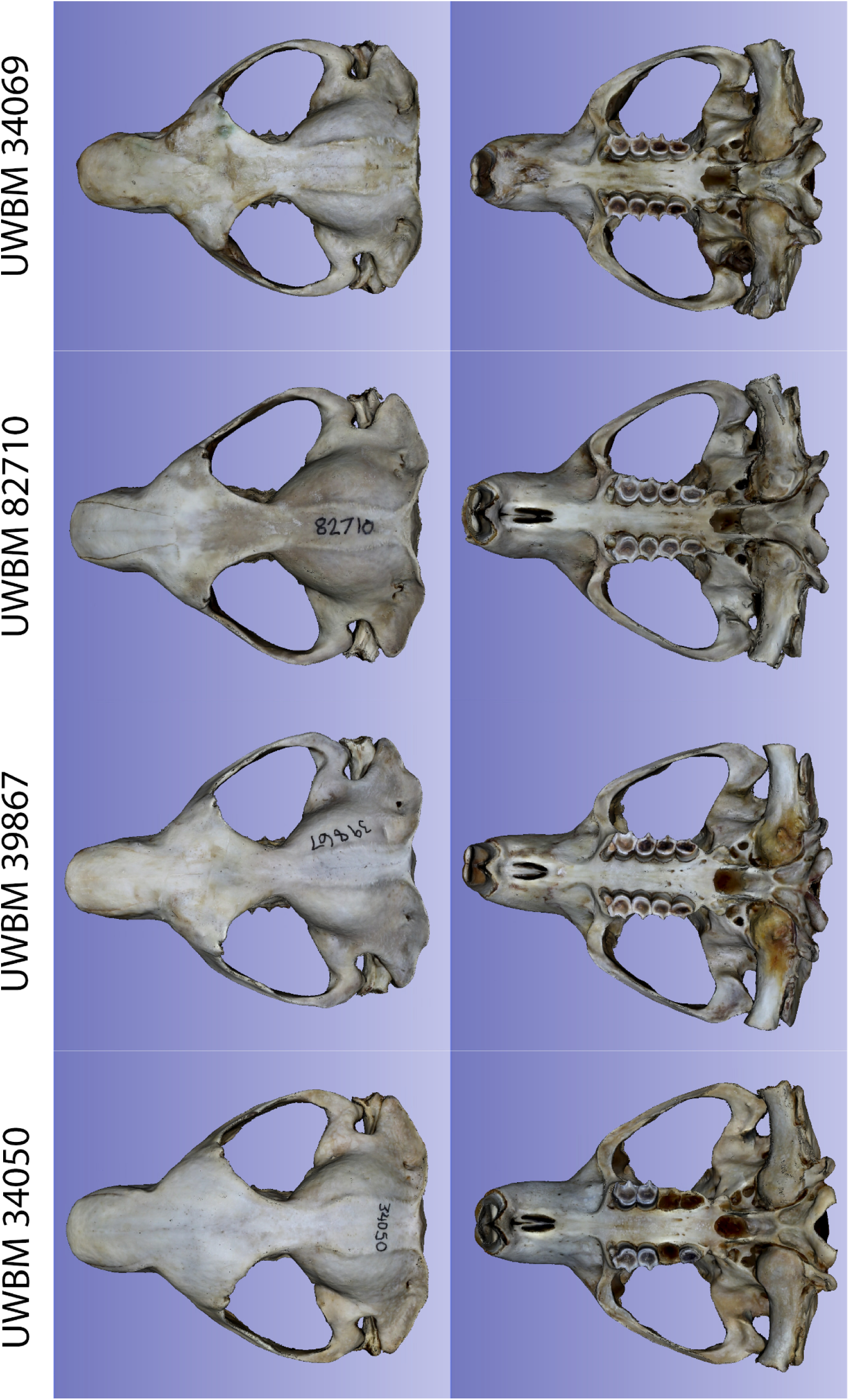
Dorsal (left columns) and ventral (right columns) views of four mountain beaver (*Aplodontia rufa*) skull specimens reconstructed using the Photogrammetry extension described in this study. Specimen IDs from top to bottom: UWBM 34069, UWBM 82710, UWBM 39867, and UWBM 34050. The reconstructed models demonstrate minimal artifacts and well-defined cranial features, particularly around delicate anatomical structures such as the zygomatic arches, orbital margins, and palatal foramina, highlighting the fidelity achievable with the streamlined, open-source photogrammetry workflow.

### 3.3. Metrics and Their Interpretations

To characterize each reconstruction’s geometric accuracy, we assessed four principal error metrics—average (mean) distance from E → G, RMSE, Hausdorff distance, and Chamfer distance (Table 1). Mean distance and RMSE collectively illustrate the model’s overall fidelity. A low mean distance (∼0.32 mm) can confirm that smaller external details (zygomatic arches, sutures) are faithfully captured, assuming this value remains well within the natural morphological variation for a skull measuring 30–40 mm in length. RMSE places extra weight on larger discrepancies—thus, if we see a rise in RMSE (e.g., from 0.40 mm to 0.50 mm), it indicates that a small region of the skull may be poorly reconstructed, even if the rest is accurate. The presence of “patched” surfaces in highly fenestrated bones underscores the need for thorough photo coverage.

Hausdorff distance identifies the worst-case discrepancy—often localized around missed camera angles or reflective edges. While a moderate rise in Hausdorff suggests incomplete imaging, the effect on ecological interpretations is less dire when the overall mean and RMSE are substantially reduced. Chamfer distance, a bidirectional average, usually aligns with mean error but also captures unmatched geometry. In our case, internal cranial structures present in CT but absent in photogrammetric data inflate G → E (CT → external surface) errors, highlighting the intrinsic mismatch of internal vs. external anatomy. Researchers focusing on external shape typically emphasize E → G, ensuring a more direct measure of how faithfully the external model reproduces the micro-CT’s outer surface.

### 3.4. Utility for Ecological and Evolutionary Applications

Our approach supports a wide range of morphological studies by reducing reconstruction errors in structurally complex regions. For instance, subtle shape differences on the mountain beaver skull—such as small alveolar expansions, ridge formation, or cranial curvature—become more readily detectable. This allows researchers to investigate diet-induced shape variation, population divergence, or interspecific comparisons without extensive manual correction. High-fidelity models also facilitate geometric morphometric analyses (e.g., dense or semi-landmarks) that rely on consistent surface detail.

Furthermore, museum collections benefit from streamlined 3D digitization workflows. Minimizing post processing steps lets curators digitize large collections of specimens more rapidly, expanding digital archives of rare or fragile taxa. Such archives aid conservation, teaching, and collaborative data sharing, especially for species like *A. rufa*, where direct specimen access may be restricted. Detailed external scans can also serve as stable references for future morphological or biomechanical modeling research.

### 3.5. Recommended Best Practices

Though robust software components (e.g., SAM, ODM) mitigate many typical photogrammetry hurdles, high-quality photography remains crucial. We strongly advise:

○ Dense Angular Overlap: Acquiring 60–80% overlap and multiple vertical angles ensures all structures appear in at least two images. Missing angles or rapid camera movement often cause “holes” or bridging artifacts.
○ Uniform, Diffuse Lighting: Glare and shadows degrade feature matching. A lightbox or diffuse multi-light setup typically yields clearer, more consistent images.
○ Stable, High-Resolution Camera: A tripod or remote shutter helps avoid blur, and higher f-stops increase the depth of field. Resolution at thousands of pixels per side better preserves subtle ridges or foramina.

4. Neutral Backgrounds and Optional Markers: Simple backdrops reduce confusion for SAM-based masking. If absolute scaling is required, incorporate ArUco or similar markers for precise reference distances.

5. Pre-Processing Checks: Reviewing image sets immediately prevents uncorrectable oversights. Underexposed or blurry shots can be retaken before dismantling the setup.

## 4. Conclusions

Our findings demonstrate that integrating the Segment Anything Model with NodeODM in 3D Slicer notably improves photogrammetric reconstructions of *Aplodontia rufa* skulls compared to the prior workflow (Zhang & Maga, 2023). Automating background masking and systematically optimizing parameters yielded a 10–15% reduction in mean distance and RMSE—which is particularly beneficial for thin cranial regions. These accuracy gains minimize laborious edits and enhance micro-CT alignment, making subtle morphological signals more discernible. It should be noted that further improvement in geometric accuracy can be accomplished by taking additional photographs of the problematic regions in orientations not acquired originally and supplementing the existing dataset.

Beyond lowering error rates, the updated pipeline’s open-source design supports broader taxonomic and collection management applications. By embedding image masking, reconstruction, and morphometric tools within one ecosystem, researchers and curators can efficiently generate high-fidelity 3D models, catalyzing new studies or outreach activities in functional morphology, ecology, and museum digitization using an open-source platform.

## Supplementary Figures

Each supplemental figure (Figures S1–S11) illustrates the geometric accuracy comparison between the updated photogrammetry workflow presented in this study and the previous method from Zhang et al. (2023) for individual mountain beaver (*Aplodontia rufa*) skull specimens. Each figure depicts dorsal, ventral, and oblique posterior-lateral views of the specimen reconstructed using our Photogrammetry module, colored by distance deviation from the corresponding micro-CT ground-truth reference. Warmer colors (red shades) represent areas with higher deviation, whereas lighter colors (yellow shades) indicate regions of low deviation. Across all specimens, the updated workflow consistently reduces artifacts and improves reconstruction quality, particularly around delicate cranial structures such as orbital margins, zygomatic arches, and foramina.

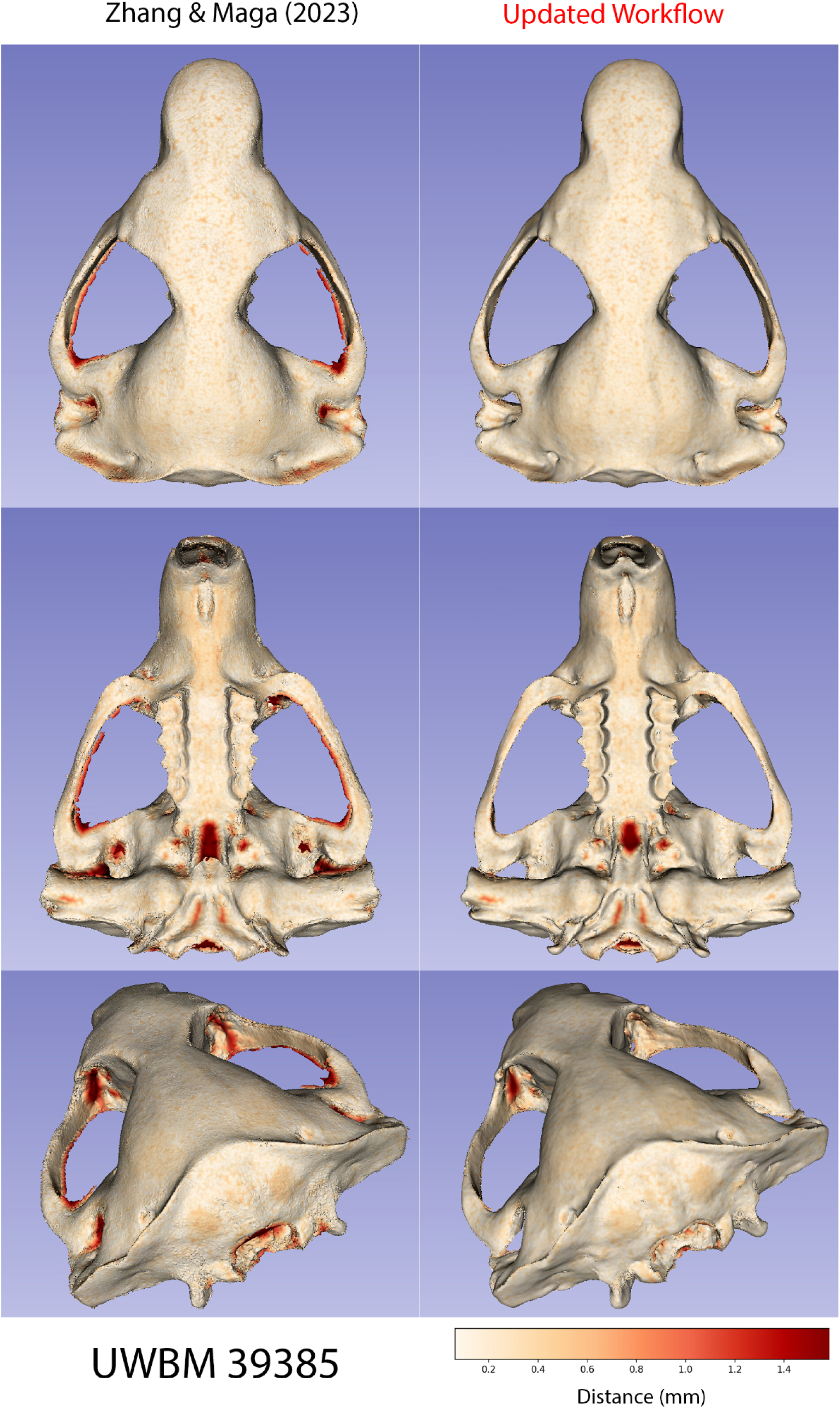

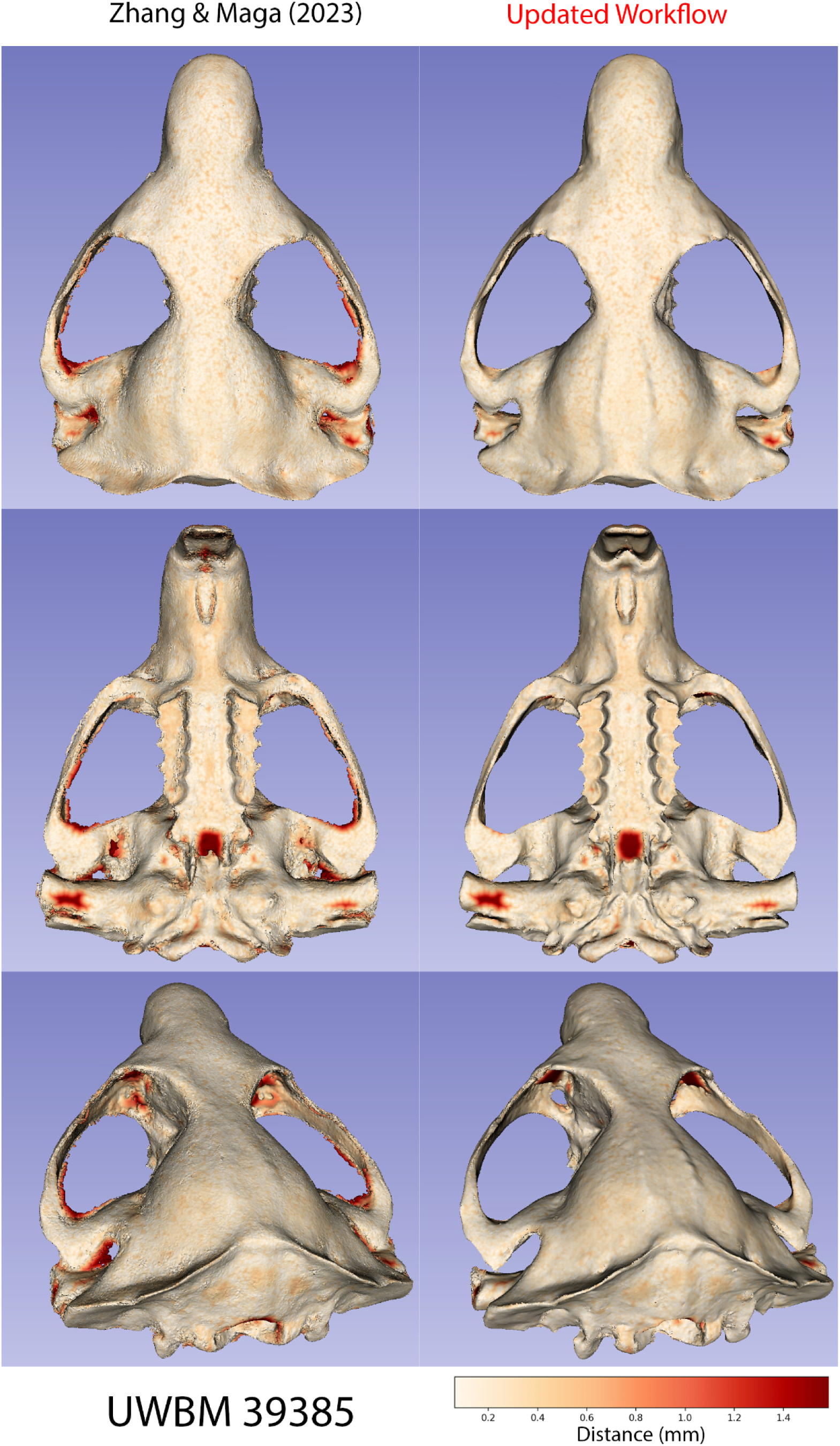

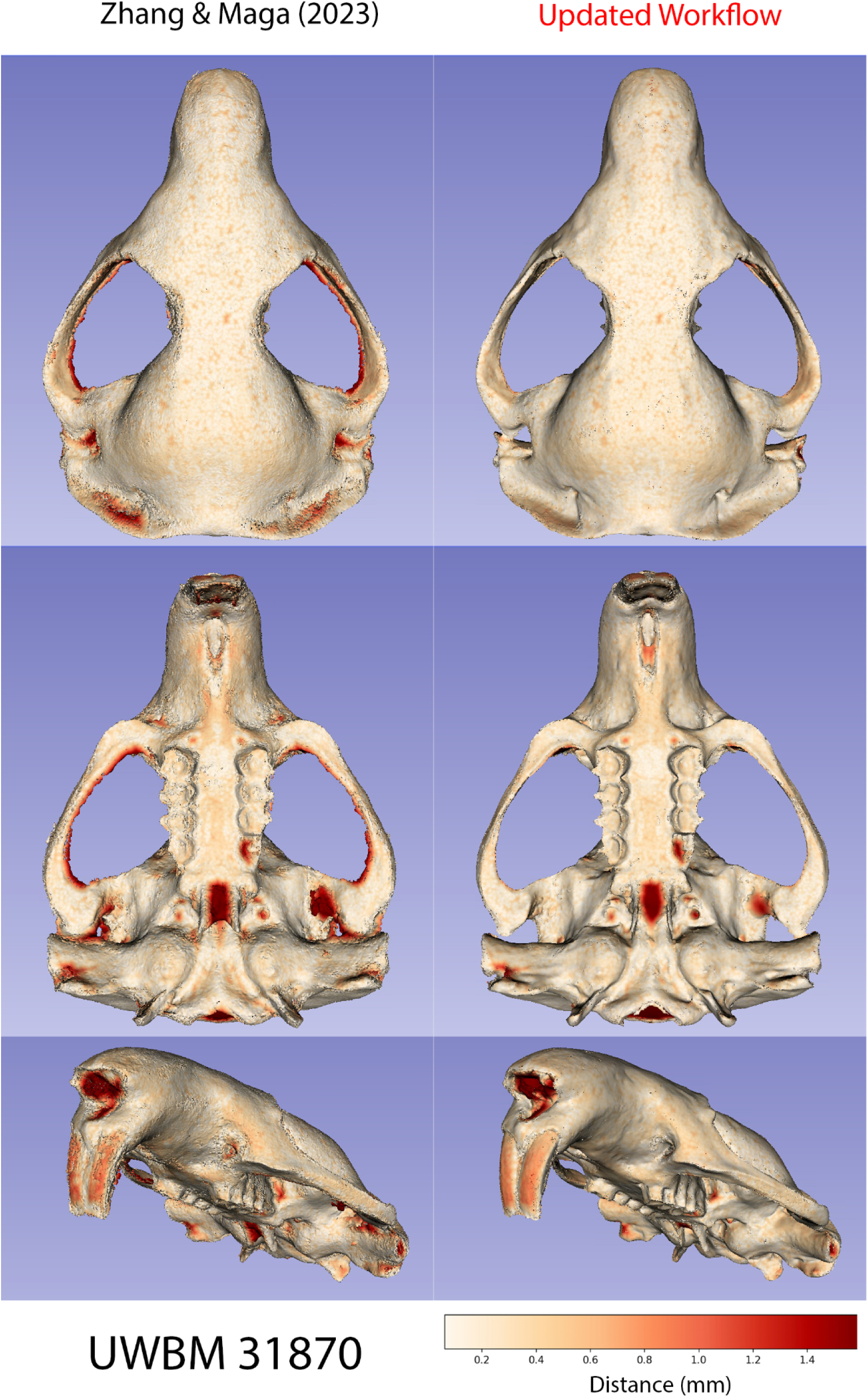

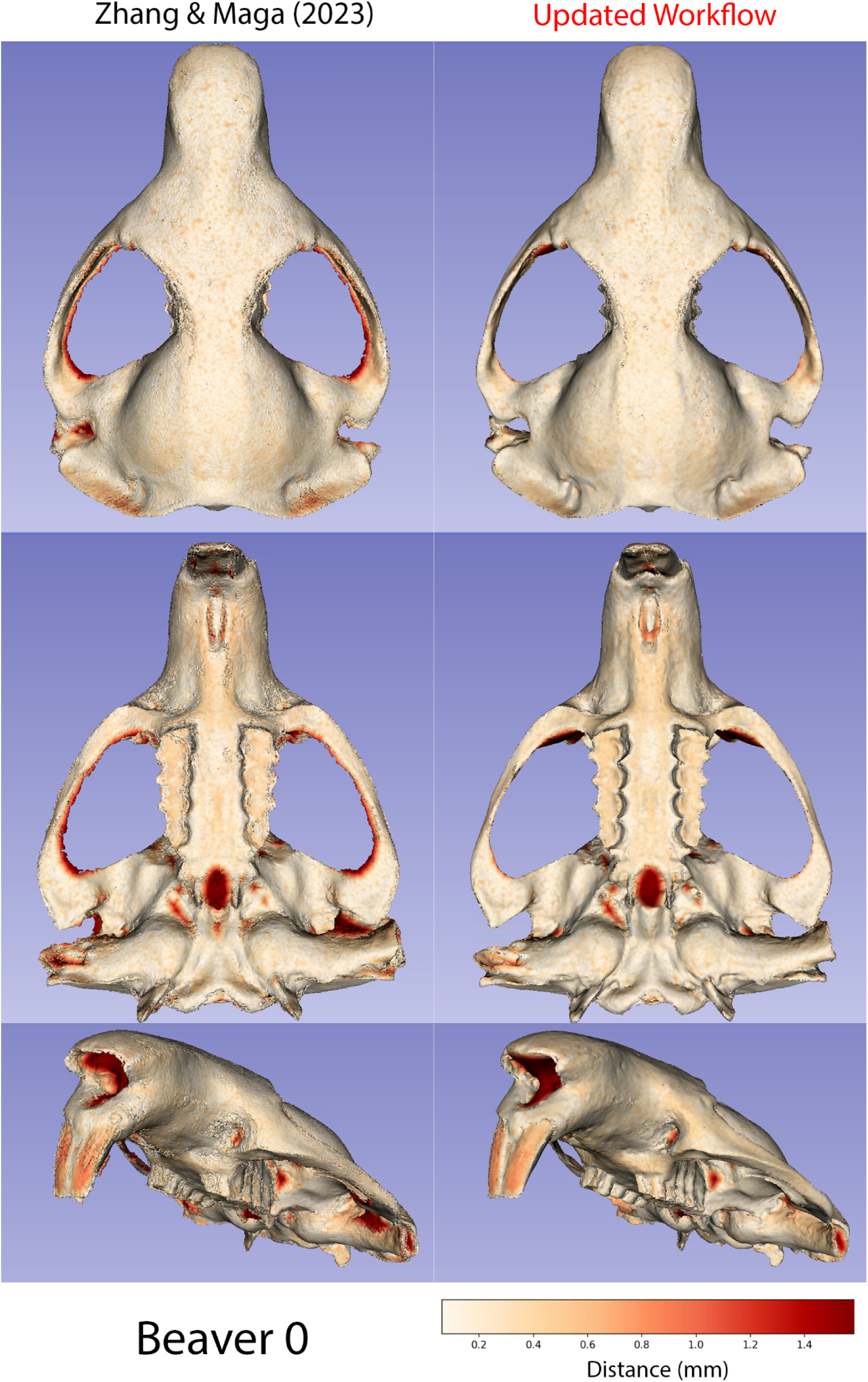

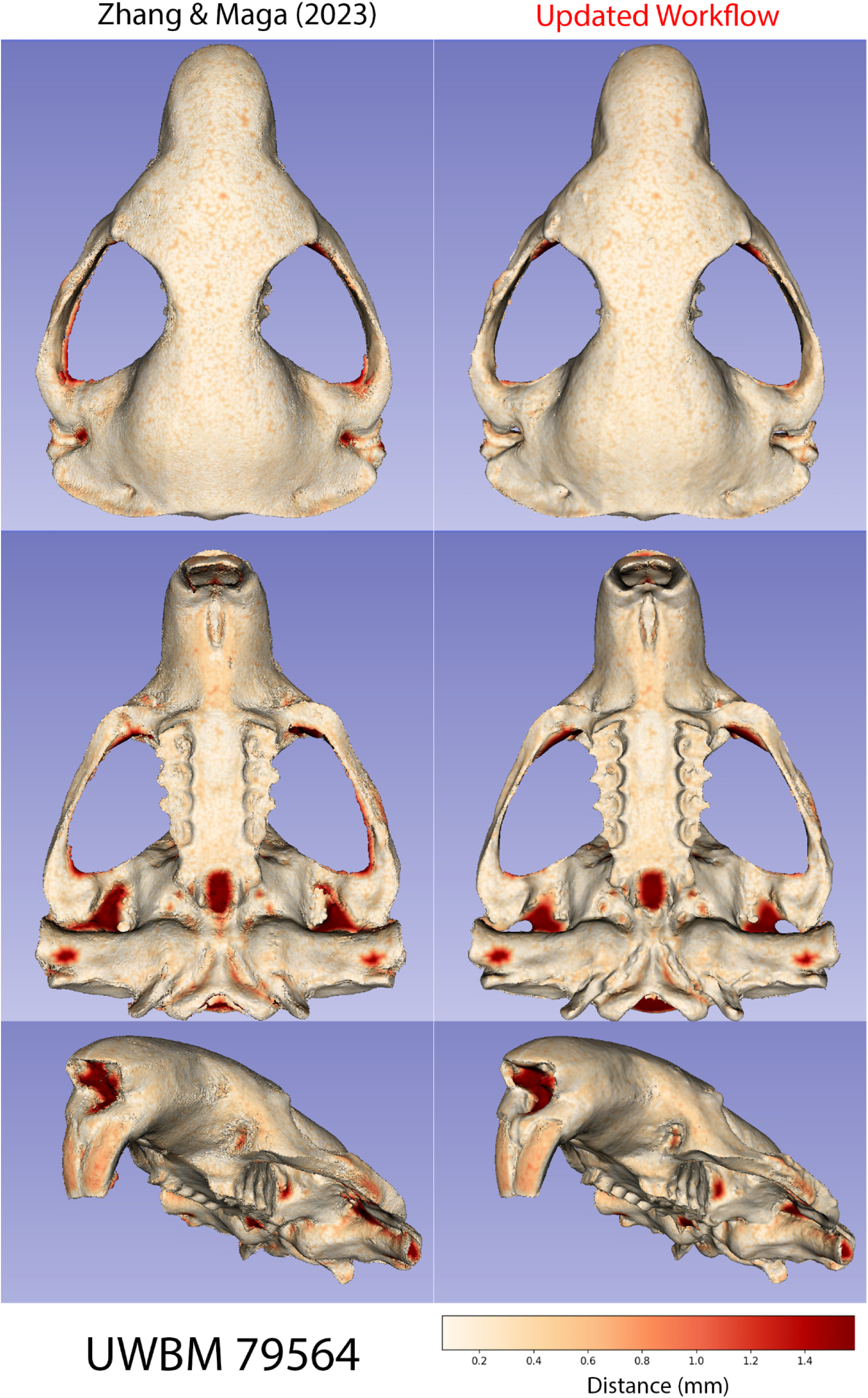

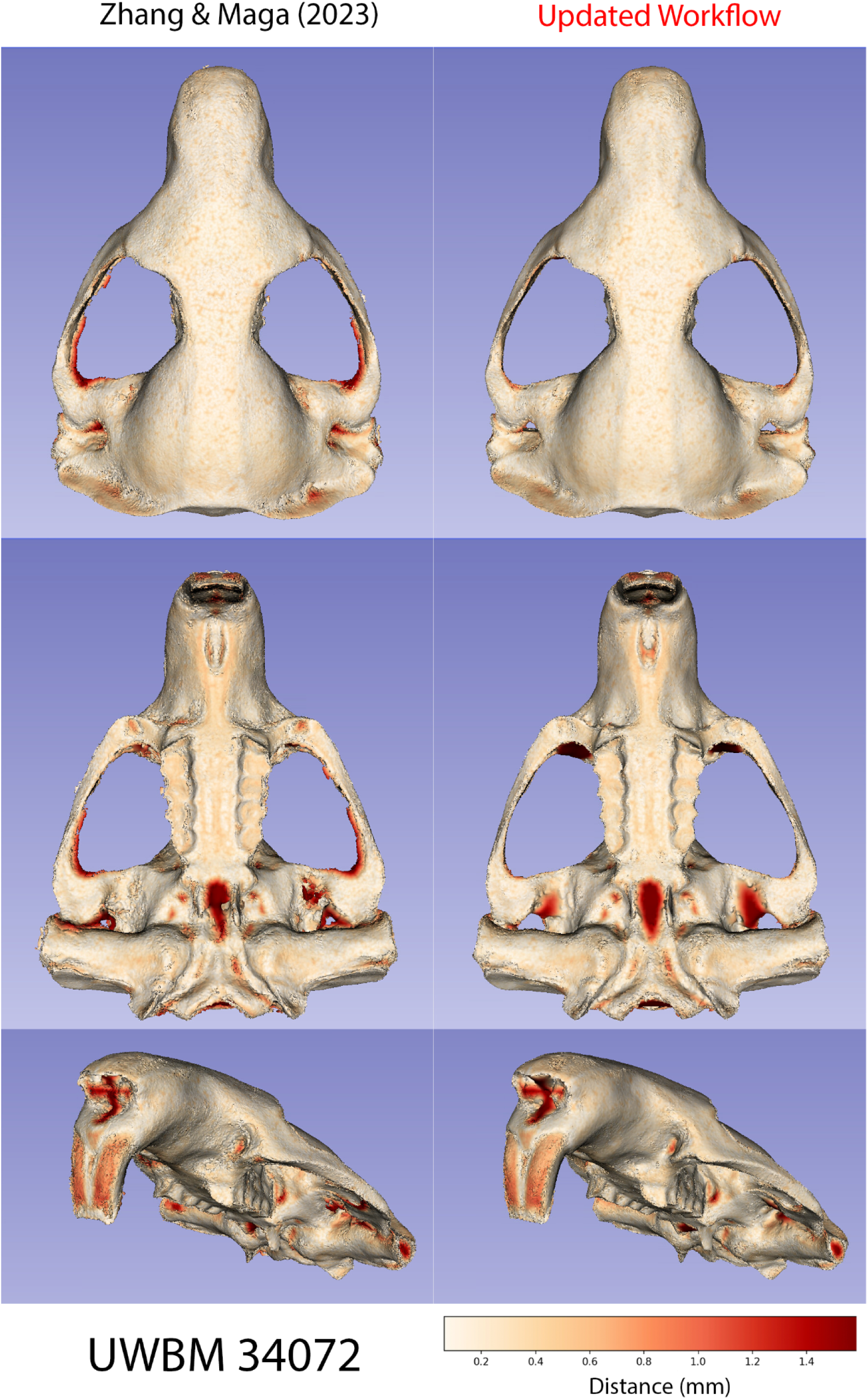

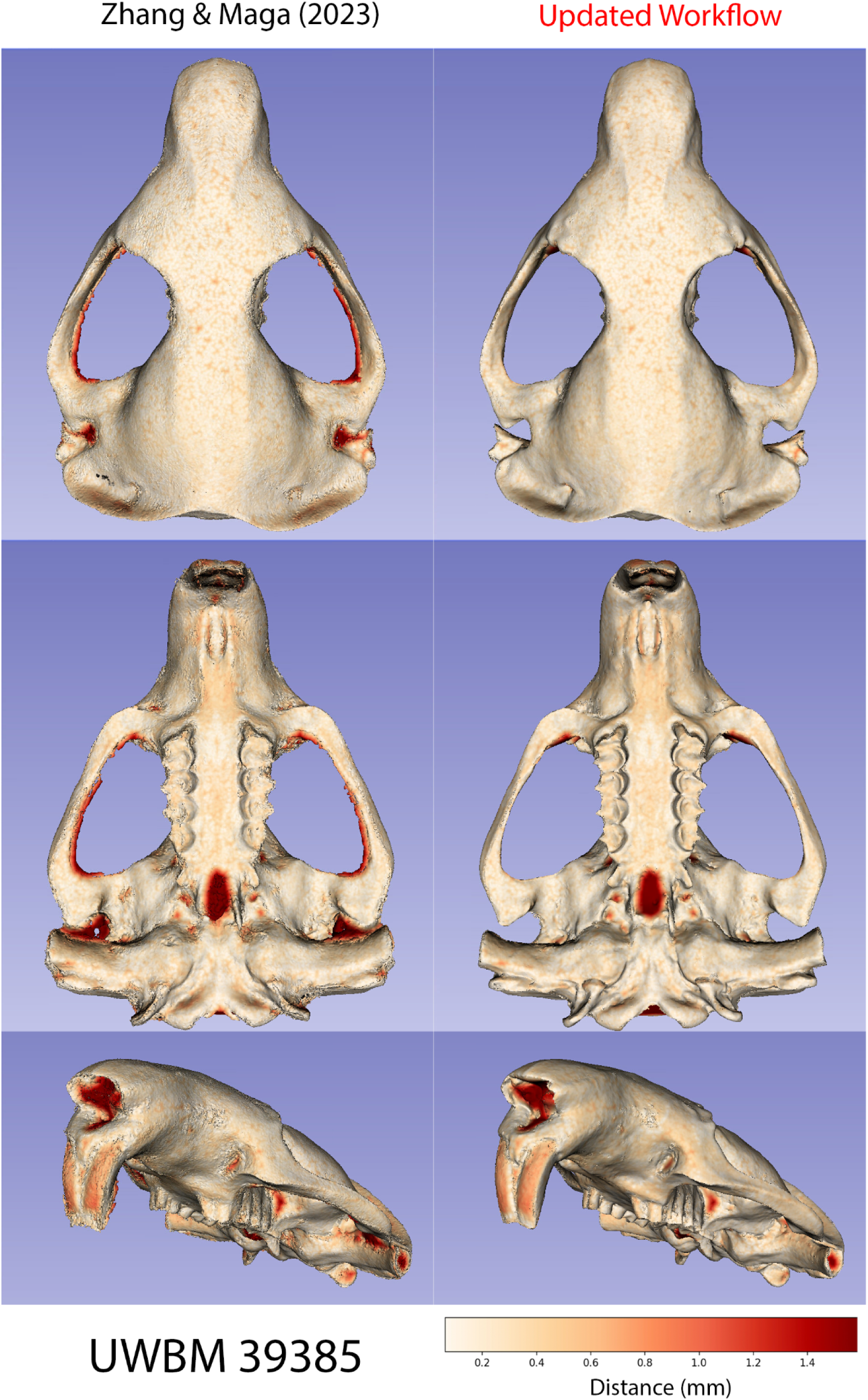

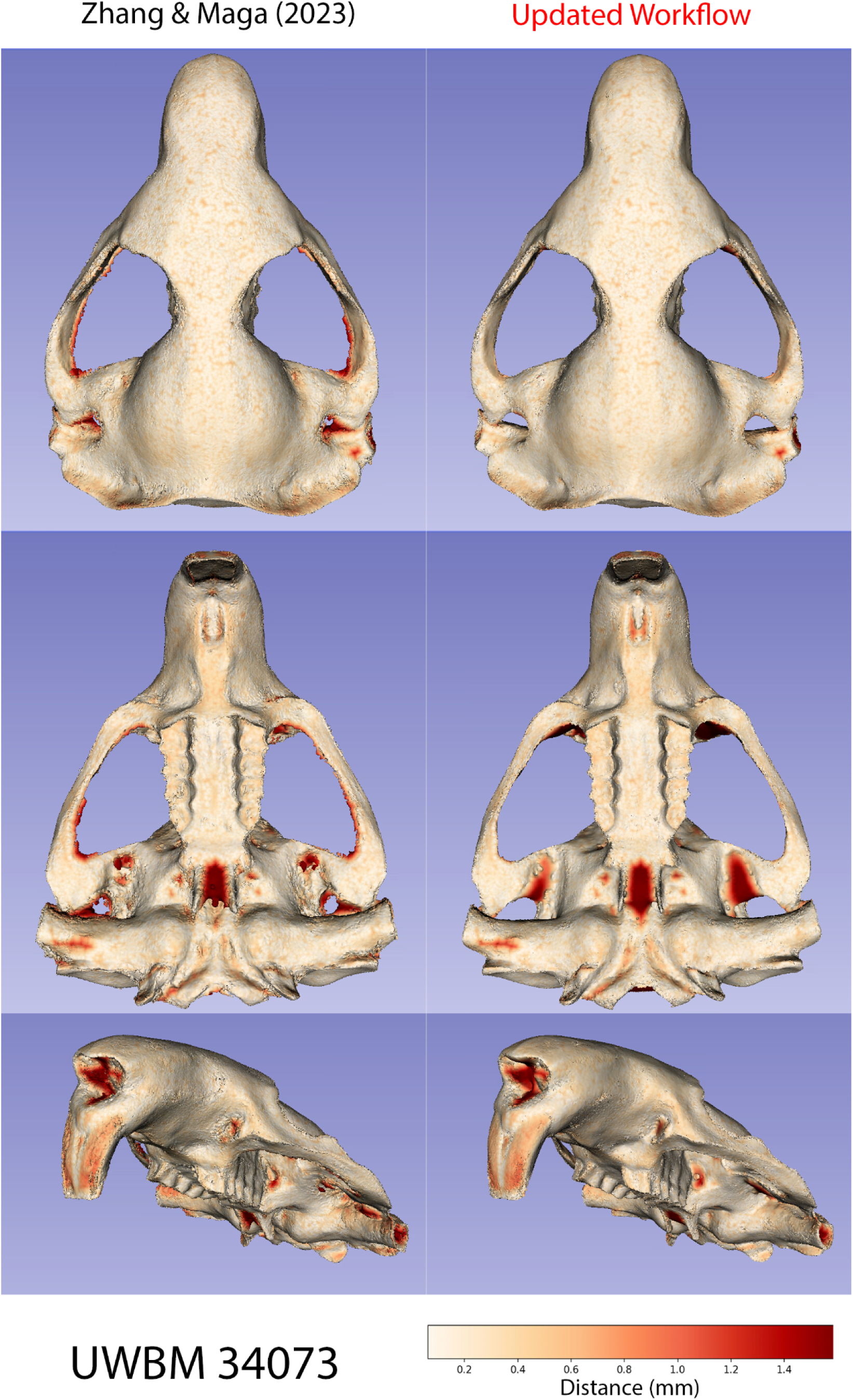

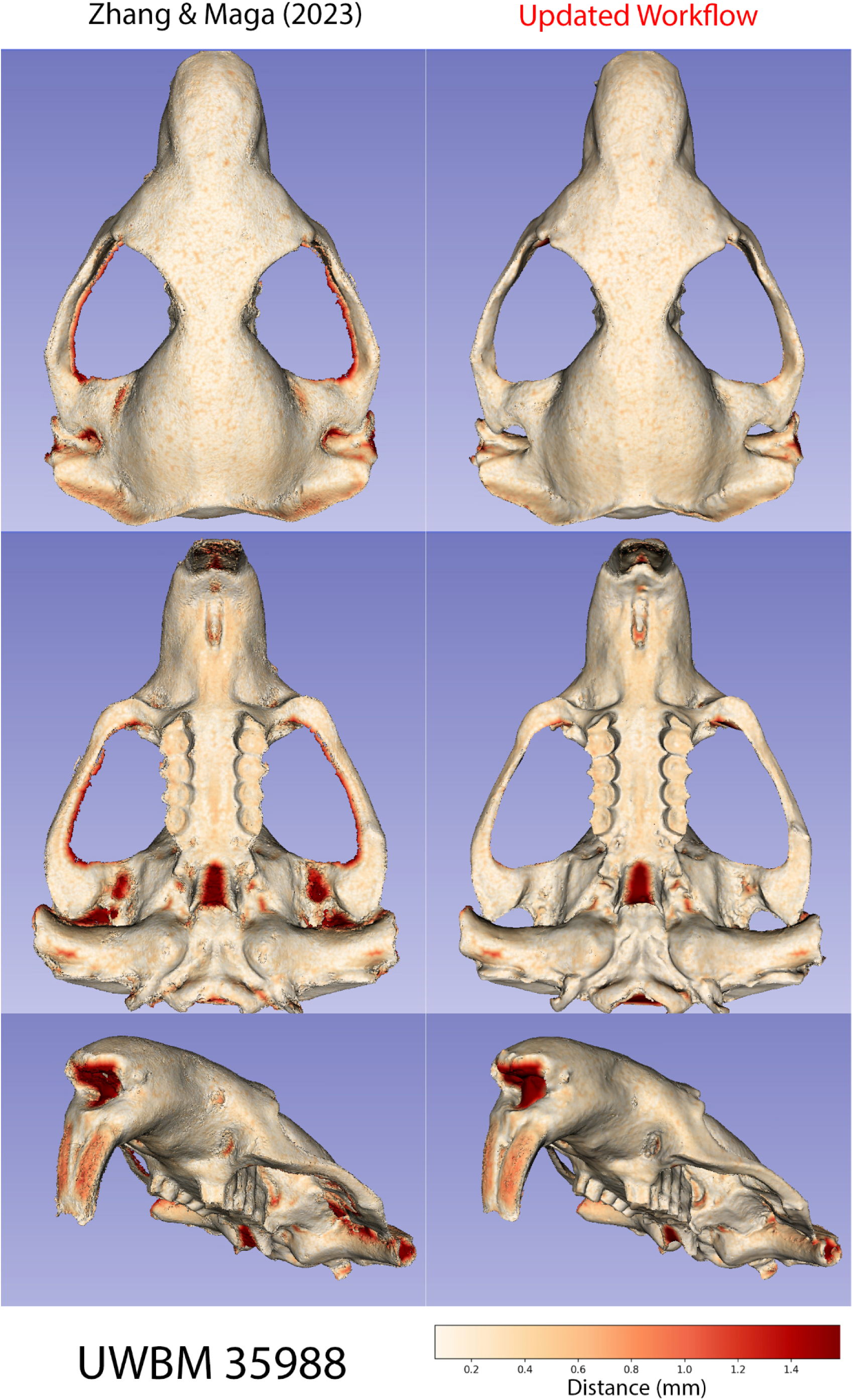

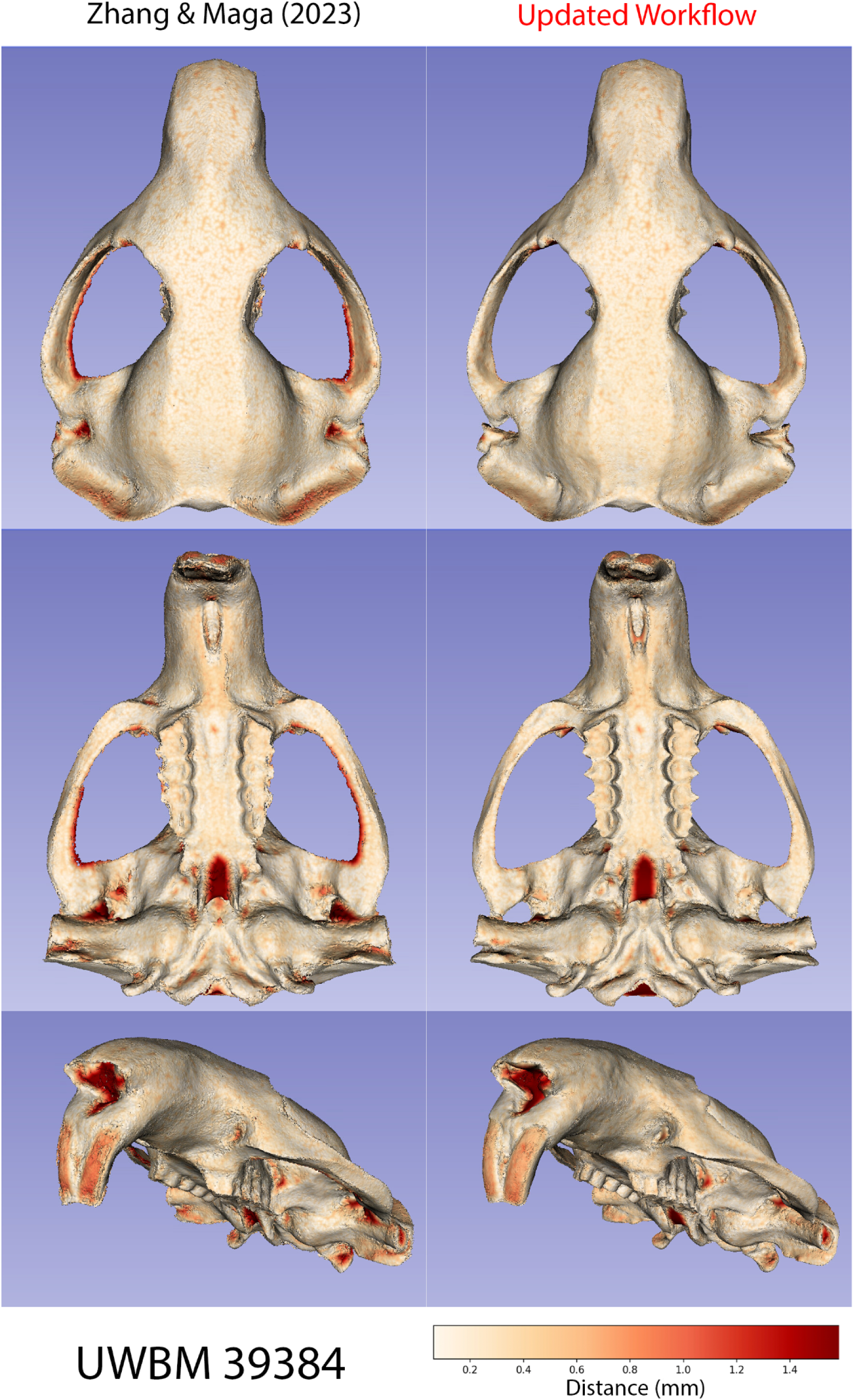

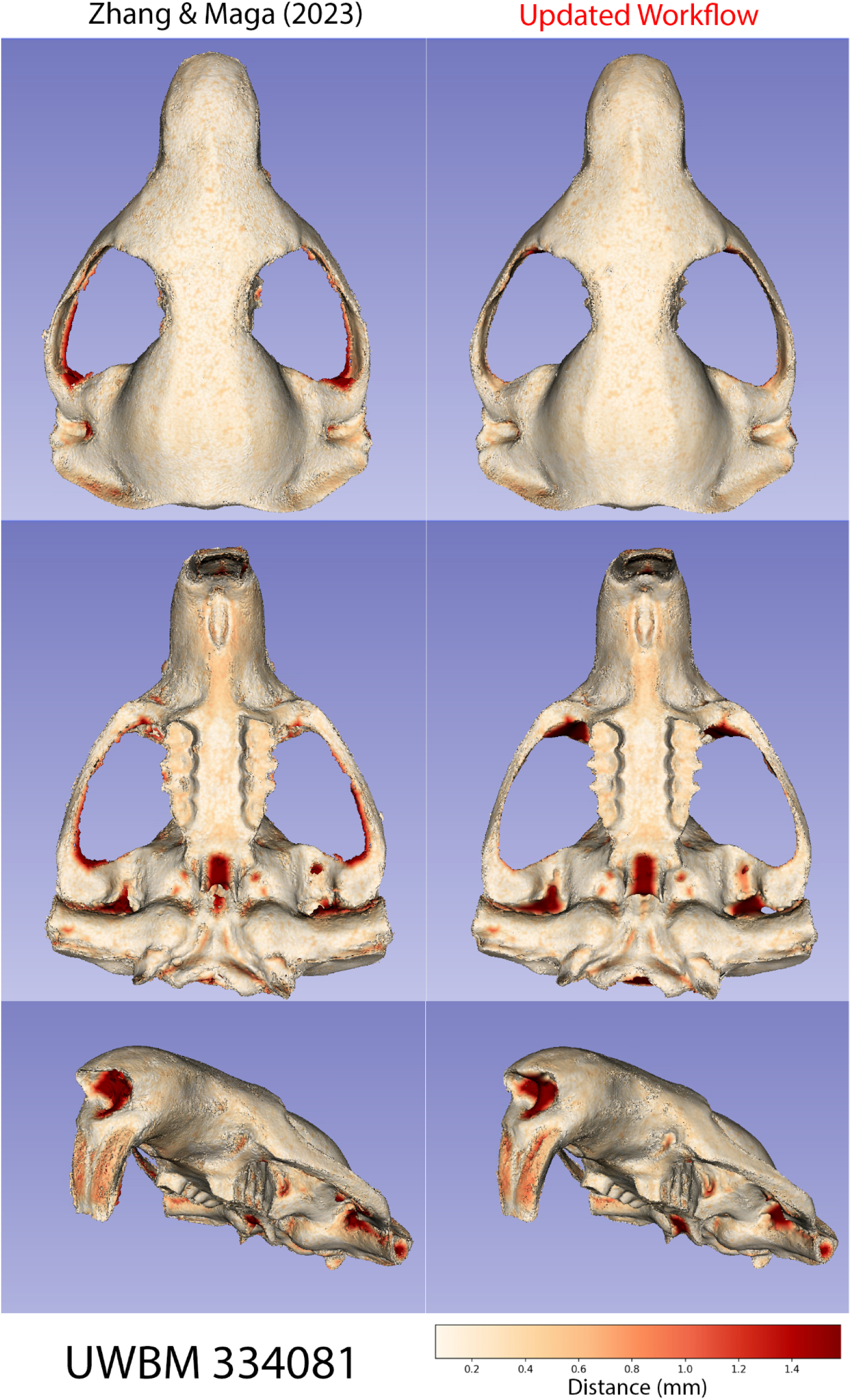

